# Self-timed movement initiation requires rapid sequential coordination of circuits in prefrontal cortex and cerebellum

**DOI:** 10.64898/2026.07.23.739940

**Authors:** Mikhail V. Monakhov, Gregory J. Wojaczynski, Javier F. Medina

## Abstract

The neural ‘chain of events’ necessary for movement initiation is poorly understood, particularly for endogenous actions that are self-timed. Here, we introduce a new task that requires mice to self-initiate a movement with high temporal precision in the absence of external ‘go’ signals. Brief, precisely timed flashes of optogenetic photoinhibition reveal that movement initiation is causally driven by rapid sequential recruitment of neurons in prefrontal cortex and lateral cerebellum.

## MAIN

To perform an action successfully, the brain must not only select *what* movement to make, but also control *when* to make it^1^. Movement initiation (MI) is under the control of a network of interacting neural circuits^2^, but we are only beginning to understand how the activity of these different brain areas is coordinated on millisecond timescale to determine the precise moment a motor response should start. Much of what we know about the brain-wide neural dynamics underlying movement initiation comes from studies in which the motor response is prepared ahead of time and triggered at the end of a delay period by an external ‘go’ cue^2,3^. For these externally driven actions, the ‘go’ cue activates reticular and pedunculopontine neurons that project via the thalamus to the motor cortex^2^, leading to the recruitment of subcortical brain regions^3,4^, and causing a rapid transition from planning-related activity to the motor command responsible for initiating movement^3^. In contrast, we know less about the multi-regional cascade of neural activity leading to self-initiated movements in the absence of external ‘go’ cues. There is general agreement that cortico-basal ganglia networks play a key role in initiating these internally driven actions (for reviews, see^5,6^). However, previous work suggests that cortico-cerebellar circuits may be engaged as well^1,7,8^, and may contribute to initiation^9–12^, especially when the start time of the motor response must be controlled with millisecond precision without the aid of external cues^13–16^.

To investigate the causal role of cortico-cerebellar pathways in the initiation of self-timed movements, we developed a new behavioral task for mice – the omitted oddball (OO) task – based on similar tasks designed to study the neural basis of internal timing in humans and monkeys^17,18^. In the OO task (**Fig.1a**), mice learn to generate an endogenous ‘go’ signal that triggers an anticipatory protective eyeblink just before the delivery of an aversive eyepuff. Unlike in classical eyeblink conditioning tasks with external sensory ‘go’ cues, the ‘go’ signal in the OO task must be generated internally by dynamically updating a prediction about the next occurrence of a periodically repeating LED flash stimulus and detecting when that next stimulus has been omitted (**Fig. 1a, right**). Mice were able to learn this challenging task, even though it took considerably longer compared to similar tasks in which the impending delivery of the eyepuff is overtly signaled with an external ‘go’ cue (**Extended Data Fig. 1**). To rule out the possibility that mice simply learned to wait until the offset of the last LED stimulus and responded at a fixed latency after that, we trained other mice in two control tasks: a standard trace conditioning task with a single LED flash, and a “trace offset” task where the train of LED flashes was replaced with one continuous long light pulse (**Extended Data Fig. 1a**). The probability of initiating the blink movement in these two control tasks was highest 100-200 ms after the offset of the LED flash, whereas in the OO task it was 400-500 ms after the offset of the last LED flash (corresponding to 100-200 ms after the time of the ‘omission’) (**Extended Data Fig. 1a**). Furthermore, when mice were trained with other intervals between LED flashes, the latency of the movements varied predictably relative to the last LED flash but remained time-locked relative to the ‘omission’ (**Fig. 1b-e**, **Extended Data Fig. 2**). These results indicate that in the OO task, mice rely on internal predictions to detect the precise time of the omission and initiate the protective eyelid movement.

**Fig. 1.**
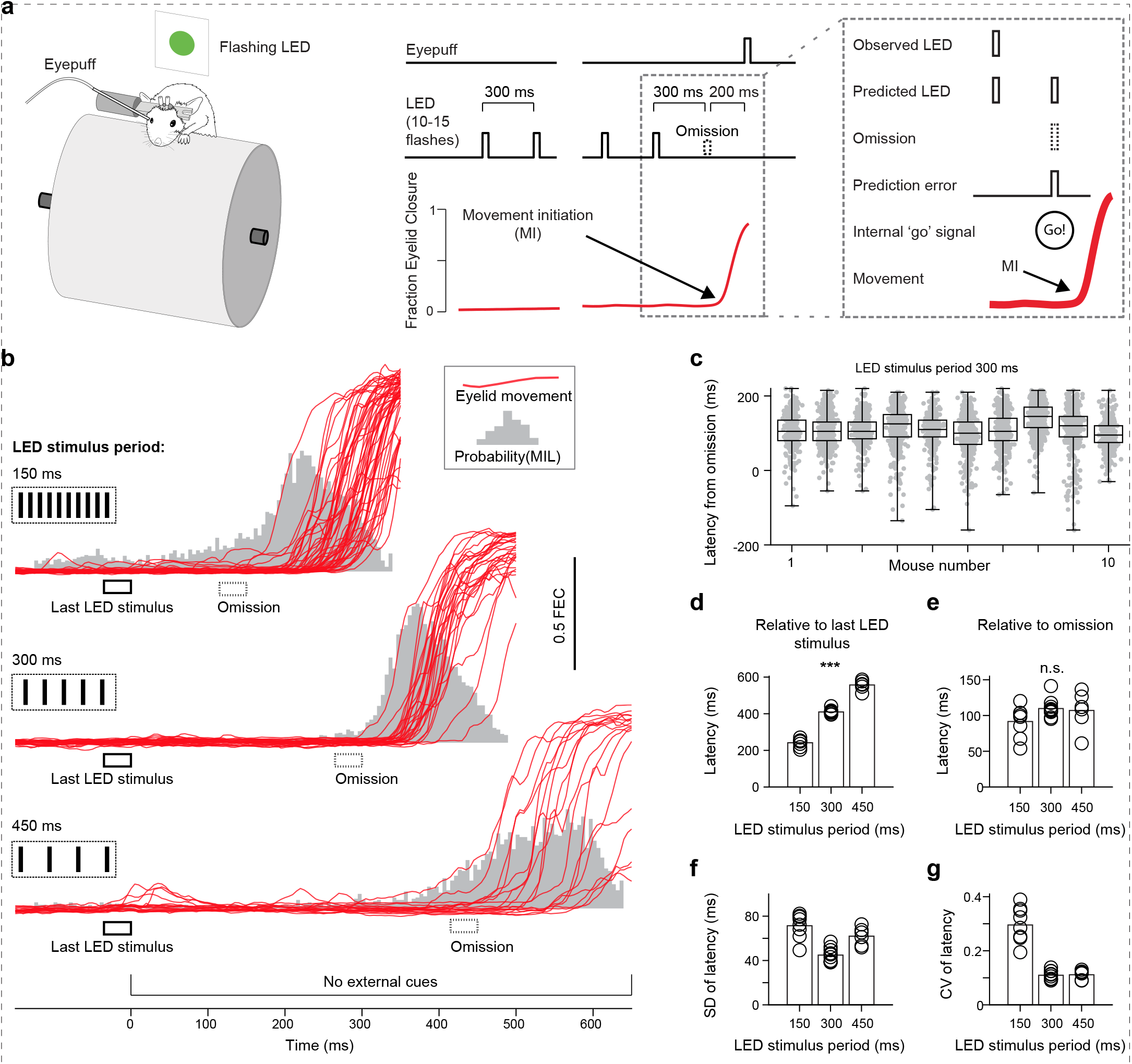
A new task to study self-timed movements with high temporal precision in mice. **a,** The mouse is head-fixed on top of a freely rotating cylinder. Green LED flash is used as periodic isochro-nous stimulus (random number of flashes, 10 to 15). Aversive eyepuff is directed to the right eye. The eye is recorded with high-speed camera to measure eyelid movements. An internal ‘go’ signal to initiate movement is generated at the time of stimulus omission, when the next predicted stimulus is not observed. MI, movement initiation. **b,** Examples of self-timed eyelid movements in three mice trained with different LED stimulus periods. MIL, movement initiation latency. FEC, fraction eyelid closure. **c,** Movement initiation latencies in mice trained with 300 ms LED stimulus period (n = 10 animals). The latencies are calculated relative to the moment of LED stimulus omission. Each marker shows movement onset in one trial. Error bars, Mean ±SD. **d,** Average movement initiation latencies relative to the last LED stimulus, in the 3 groups of mice trained with different values of LED stimulus period (150 ms, 300 ms and 450 ms; n = 8, 10 and 6 mice respective-ly). Markers show averages across trials for each mouse. Bars are averages across mice. *** One-way ANOVA, F(2, 21) = 444.52, P < 0.001. **e,** Same as in **(d)**, but latencies are calculated relative to the moment of LED stimulus omission. n.s., not statistically significant. One-way ANOVA, F(2, 21) = 2.04, P = 0.155. **f,** Standard deviations of movement initiation latencies. **g,** Coefficients of variation of movement initiation latencies (relative to last LED stimulus).

We assessed the contribution of cerebellum and prefrontal cortex to movement initiation in the OO task, using optogenetic tools to inhibit neurons in these brain areas for a very brief moment around the time of the omitted LED stimulus. These experiments are technically challenging and would not have been possible without the high temporal resolution of optogenetics and the remarkably consistent movement latencies we observed in the OO task, especially in mice trained with the 300 ms LED-LED interval (**Fig. 1f-g**). In addition, it was necessary to train the mice in the presence of a randomly intermittent flashing blue light (**Extended Data Fig. 3**), to mask the laser light and prevent it from interfering with detection of the omitted LED stimulus (**Extended Data Fig. 4**). Although the intermittent light caused a slight drop in performance (**Extended Data Fig. 3b,c**), mice were still able to initiate the movement successfully even when the laser light was flashed outside the brain at the time of the omitted LED stimulus (**Extended Data Fig. 3d-f**). These controls rule out the possibility that any performance deficits we may observe in the optogenetic experiments described below may be simply caused by interference from the laser light itself.

First, we examined how mice performed in the OO task during optogenetic inhibition of neurons in the cerebellar dentate nucleus (DN), a brain area necessary for initiating movements with high temporal precision in other tasks^9,12,16,17^. We used transgenic mice with channelrhodopsin in inhibitory Purkinje cells and stimulated their axon terminals to suppress DN activity (**Fig. 2a, Extended Data Fig. 5**). Because our objective was to define the exact time at which DN makes its contribution to movement initiation, we used very brief laser pulses (35 ms ‘opto-blink’) and varied their timing relative to the omitted LED stimulus (**Fig. 2b**). Opto-blinks of the ipsilateral DN (same side as the eye receiving the puff) in the first 100 ms after the omitted stimulus caused a delay in movement initiation that reached its maximum when the photoinhibition started 70 ms after the omission (**Fig. 2c-j**). Further analysis and control experiments confirmed that this delay was caused by transiently preventing movement initiation during a 65-75 ms window after the onset of the opto-blink (**Extended Data Fig. 6**), and could not be simply explained by post-inhibitory rebound excitation of DN neurons^19^ (**Fig. 2g** shows no photoinhibition-driven movements when the laser pulse was delivered by itself in non-trial periods), or by the laser light (**Fig. 2c-e** shows movements were not delayed when the contralateral DN was photoinhibited). These results demonstrate that activation of DN neurons after the omission of the LED stimulus is necessary for initiating movement in the OO task. However, because opto-blinks caused a delay but did not abolish movements altogether, we hypothesized that the internal ‘go’ signal that drives movement initiation persists during inhibition of DN and must be generated elsewhere in the brain. The dorsomedial prefrontal cortex (dmPFC) is well-positioned to play this role because frontoparietal networks including the supplementary motor area respond to unexpected stimulus omissions^20^, and because the dmPFC is a key source of persistent inputs to the cerebellum in other learning tasks^21^.

**Fig. 2.**
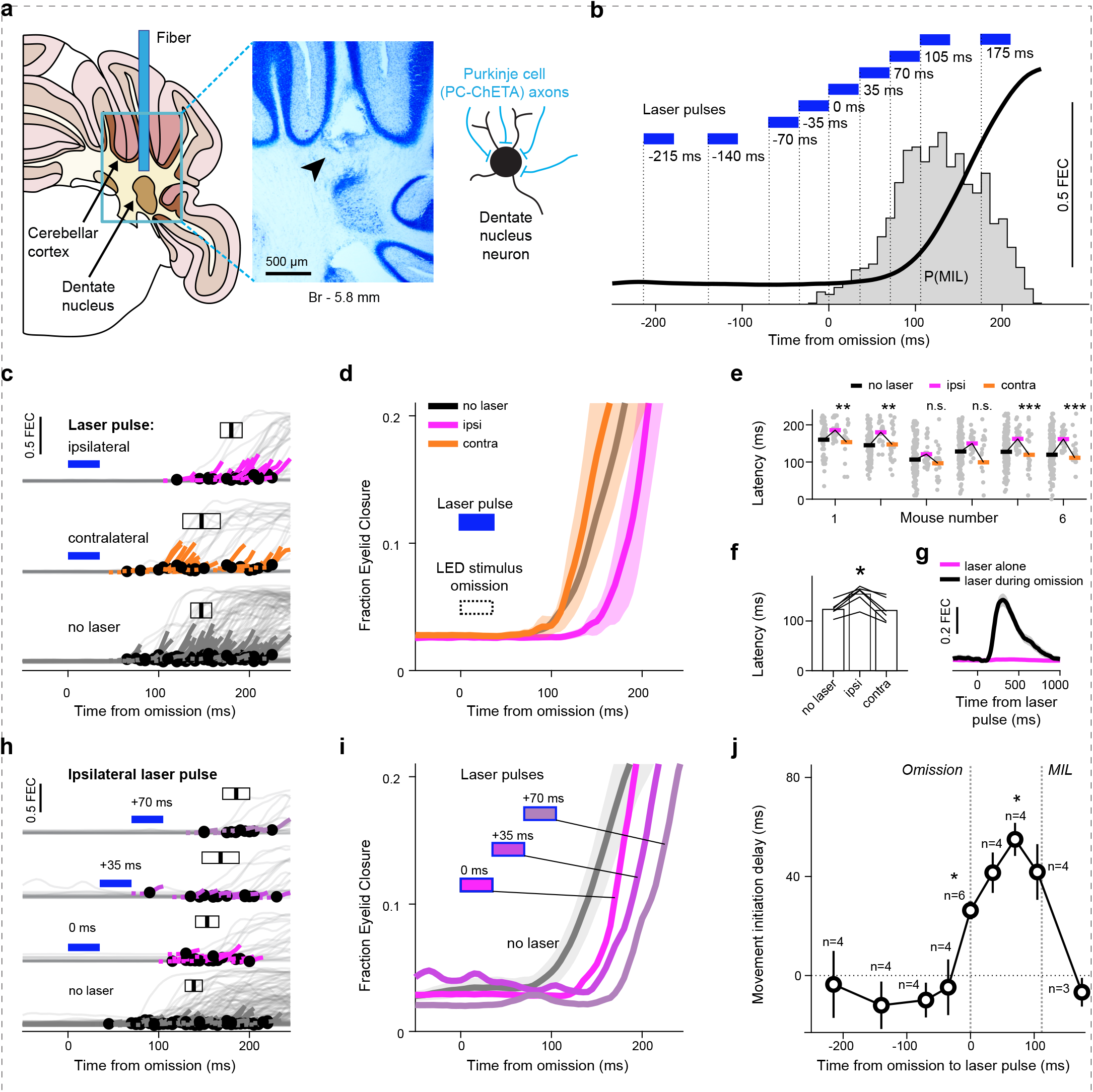
Movement initiation is delayed by brief inhibition of cerebellar dentate nucleus. **a,** The position of optic fiber in cerebel-lum. The arrowhead on histology section shows estimated location of fiber tip. The mouse expressed channelrhodopsin in inhibitory Purkinje cells. **b,** Timing of laser pulses used for optogenetic inhibition of dentate nucleus. Blue bars are laser pulses (only one pulse was delivered in any given trial). Thick black trace shows averaged eyelid movement (without laser). FEC, fraction of eyelid closure. The histogram in background shows probability distribution of movement initiation latencies (MIL), n = 6 mice. **c,** Inhibition of dentate at the moment of LED stimulus omission (example session). Black dots show movement onsets. Vertical bars, mean latency. Black rectangles, 95% CI. **d,** Averaged eyelid movements from the same session as in **(c)**. Shaded areas show 95% CI (only for trials with laser). **e,** Movement initiation latencies in individual mice (n = 6), in trials without laser (no laser), or with laser in the ipsilateral (ipsi, same side as the eye receiving the puff) or contralateral (contra) dentate nucleus. Data for individual trials (gray markers) and the averaged latency in each condition (colored horizontal bars). Two-sample t-tests (ipsilateral inhibition vs all other trials), significance indicated for each mouse, n.s. P > 0.05, ** P < 0.01, *** P < 0.001. **f,** Comparisons of averaged latencies (n = 6 mice). Laser was delivered at the moment of LED stimulus omission. * Ipsi vs contra, paired t-test, t(5) = 4.85, P = 0.005. Ipsi vs no laser, paired t-test, t(5) = 6.21, P = 0.002. **g,** Average eyelid movement following laser pulses delivered at the time of LED stimulus omission (black trace) or during the intertrial interval (magenta trace); n = 6 mice. **h,** Same as in **(c)**, for different laser timings (example session). Labels indicate the timing of the laser pulse relative to the moment of LED stimulus omission. **i,** Averaged eyelid movements from the session in **(h)**. **j,** Average delay in movement initiation for experiments with different laser timings. Mean ±SEM. MIL, movement initiation latency (average across mice). * P < 0.05, one-sample t-tests with Bonferroni correction (9 comparisons), n = 3 to 6 mice at each time point. Time=0 ms: t(5) = 5.32, P = 0.028; Time=70 ms: t(3) = 8.23, P = 0.034.

We used the opto-blink approach to define the role of the dmPFC in the OO task (**Fig. 3a,b**). Brief inhibition of dmPFC output neurons was achieved by optogenetically stimulating local parvalbumin interneurons (**Fig. 3a**). In striking contrast to the effects observed for the DN (**Fig. 2**), bilateral opto-blinks of the dmPFC did not change the latency of movement initiation (**Fig. 3c, Extended Data Fig. 7**) or inhibit ongoing movements (**Fig. 3d**). Instead, brief photoinhibition of the dmPFC caused a profound decrease in both the probability and the amplitude of the movement, but only when the laser pulse was given within a narrow 100 ms window before or after the time of the omitted LED stimulus (**Fig. 3e,f**). Thus, dmPFC activity is not continuously required during movement preparation or execution but becomes causally essential for initiation during a remarkably brief moment around the time when the internal ‘go’ signal is generated.

**Fig. 3.**
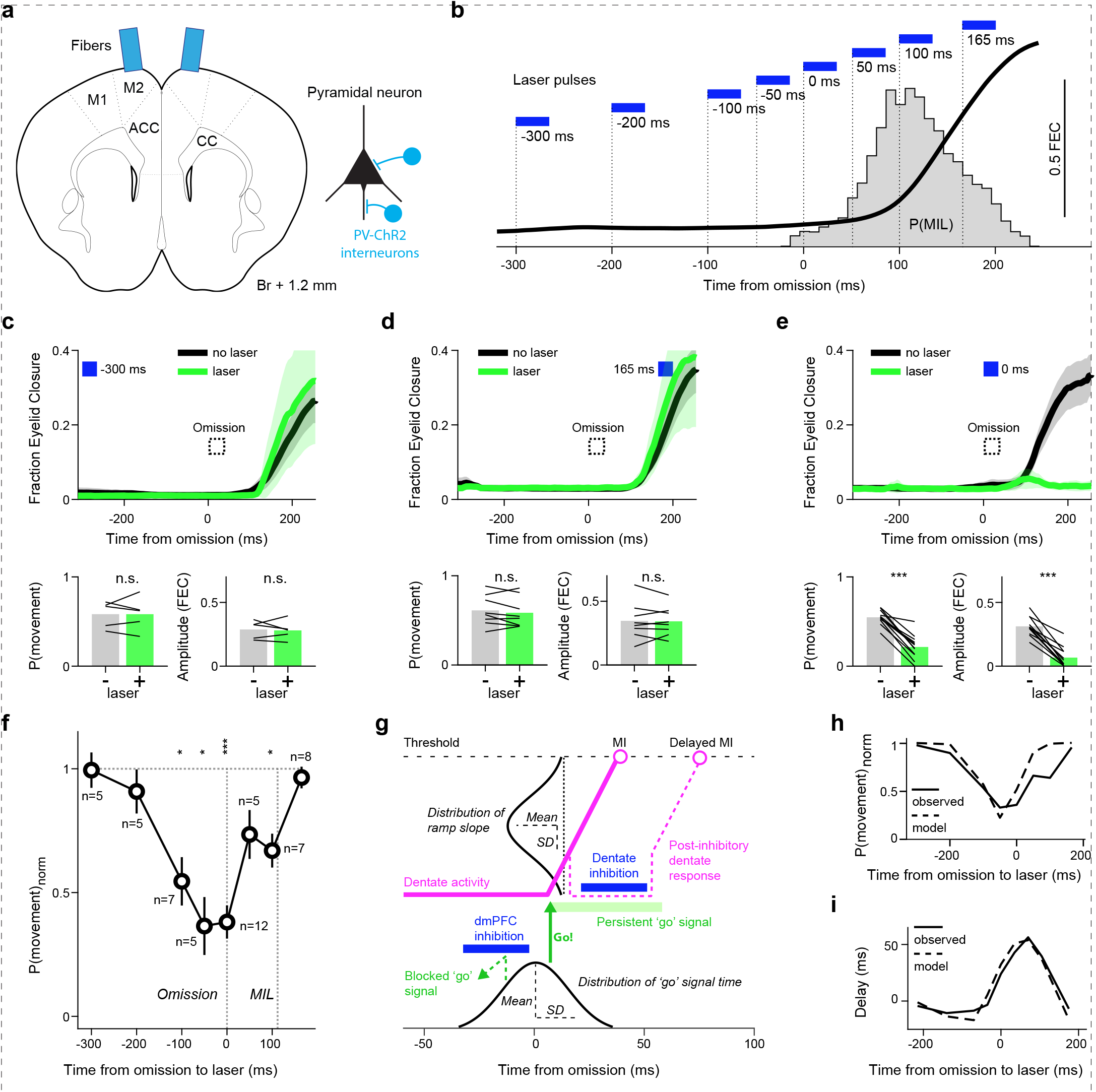
Brief inhibition of dorsomedial prefrontal cortex prevents movement initiation. **a,** The position of optic fibers above dmPFC. M1, primary motor cortex. M2, secondary motor cortex. ACC, anterior cingulate cortex. CC, corpus callosum. The mouse expressed channelrhodopsin in inhibitory parvalbumin interneurons. **b,** Timing of laser pulses used for optogenetic inhibition of dmPFC. Thick black trace is the averaged eyelid movement. The histogram in background shows distribution of movement initiation latencies. P(MIL), probability of movement initiation latency. FEC, fraction of eyelid closure. **c-e,** Three experiments with different time points of dmPFC inhibition. The plots show averaged eyelid movement traces from example sessions. Shaded areas, 95%CIs. Blue rectangle, laser pulse. Bar graphs show the performance in trials with (laser ‘+’) and without dmPFC inhibition (laser ‘-’). **c**, 300 ms before LED stimulus omission. n = 5 mice, paired t-tests, P > 0.05. **d**, 165 ms after the omission. n = 8 mice, paired t-tests, P > 0.05. **e**, during the omission. n = 12 mice, paired t-tests. P(movement): t(11) = 10.21, P < 0.001. Amplitude: t(11) = 10.61, P < 0.001. **f,** Changes of movement initiation probability after dmPFC inhibition. P(movement)_norm_ = P_with laser_ /P_without laser_. Error bars indicate SEM. MIL, average movement initiation latency. * P < 0.05, *** P < 0.001. One-sample t-tests with Bonferroni correction (8 comparisons), n = 5 to 12 mice at each time point. Time=−100 ms: t(6) = −4.65, P = 0.028; Time=−50 ms: t(4) = −5.43, P = 0.045; Time=0 ms: t(11) = − 9.25, P < 0.001; Time=100 ms: t(6) = −4.84, P = 0.023. **g,** A model for movement initiation by dmPFC and dentate nucleus. The ‘go’ signal is provided by dmPFC and drives dentate neuronal activity upwards until it reaches movement initiation (MI) threshold. Optogenetic inhibition of dmPFC blocks the ‘go’ signal. Inhibition of dentate decreases its activity but doesn’t eliminate the ‘go’ signal, which persists and causes a post-inhibition response that triggers a delayed movement. **h,** Observed and simulated effects of dmPFC inhibition at various timepoints. **i,** Observed and simulated effects of dentate inhibition at various timepoints.

Together, these results suggest a simple circuit-level model for self-timed movement initiation in the OO task. In this model, an internal ‘go’ signal is sent from the dmPFC to the lateral cerebellum at the time of the omitted stimulus, where it triggers activity in the dentate nucleus that ramps toward the threshold for movement initiation (**Fig. 3g; Extended Data Fig. 8**). This model captures the distinct effects of perturbing each node in the circuit. Brief dmPFC inhibition near the time of omission prevents movement initiation because the internal ‘go’ signal is blocked before the downstream cerebellar ramp can be engaged (**Fig. 3h**). By contrast, brief dentate inhibition after the omission does not remove the upstream drive, but instead transiently interrupts the process that converts this signal into movement, delaying initiation until dentate activity can resume its trajectory toward threshold (**Fig. 3i**). Thus, the opto-blink strategy reveals two sequential and computationally distinct steps in the initiation process: the dmPFC sends the internal ‘go’ signal, and the lateral cerebellum converts that signal into a precisely timed movement trigger.

Our findings extend recent work showing that cerebellar output can act as a context-dependent trigger for learned movements. In a cued forelimb task, activation of the dentate/interpositus–thalamocortical pathway was sufficient to initiate a conditioned movement, leading to the proposal that the cerebellum can “hold the starting gun” for action onset^9,22^. The OO task reveals a different, internally timed version of this principle: when no external ‘go’ cue is available, movement initiation depends on a brief dmPFC signal followed tens of milliseconds later by cerebellar output. This does not imply that basal ganglia circuits are dispensable. Indeed, striatal and cerebellar circuits make distinct contributions to self-timing^23^, and recent work shows that cerebellar output can rapidly modulate nigrostriatal dopamine signaling through direct projections to the substantia nigra pars compacta^24^, raising the possibility that cortico-cerebellar and cortico-basal ganglia pathways cooperate during endogenous action. What may make the OO task especially dependent on cortico-cerebellar circuits is that it requires more than self-initiation: animals must learn the tempo of a repeating stimulus, predict when the next event should occur, and act when that prediction is violated. Consistent with this view, primate studies of omission detection and internalized rhythms show that cerebellar nuclear activity preferentially represents temporal prediction of periodic sensory events, whereas striatal activity is more closely linked to motor preparation^16,25,26^. Thus, our findings suggest that cortico-cerebellar circuits may be especially critical when the brain must transform a temporal prediction error into a precisely timed, self-initiated action.

## ONLINE METHODS

### Animals

Experiments were performed on adult male C57BL/6J mice. Wild-type mice were purchased from Jackson Laboratories (No. 000664). Genetically modified mice were bred in house. PV-IRES-Cre x Ai32 mice were produced by crossing B6 Pvalb-IRES-Cre (No. 017320) and Ai32 (No. 024109). Most B6 Pvalb-IRES-Cre and Ai32 mice used for breeding were a gift from Dr Nuo Li. Some of the Ai32 mice were a gift from Dr Mingshan Xue. L7-ChETA mice were produced by crossing L7Cre-2 (No. 004146) and R26-CAG-LSL-2XChETA-tdTomato (No. 017455). Both L7Cre-2 and R26-CAG-LSL-2XChETA-tdTomato mice were bred in house. All mice were at least 9 weeks old on the day of surgery and were naïve prior to the experiments. Mice were housed in an official Baylor College of Medicine climate-controlled animal facility on an inverted light/dark cycle (7 AM lights off/7 PM lights on). Experiments were performed during the dark cycle. Procedures were performed in accordance with protocols approved by the Baylor College of Medicine Animal Care and Use Committee based on guidelines of the National Institutes of Health.

### Surgery

Prior to all experiments, stereotaxic surgery was performed to implant a headplate and optic fibers, or headplate alone. Mice were anesthetized with isoflurane (1.5–2% by volume in O_2_; SurgiVet) and kept on a heating pad to maintain body temperature. In addition, Meloxicam was given perioperatively to reduce swelling and provide post-operative analgesia. A midline incision was made to expose the skull and the underlying fascia was cleared with cotton swabs. Two small screws were placed on either side of the midline near bregma. A thin stainless steel headplate was then placed over bregma such that the screws fit into the central hole in the headplate and secured to the screws and skull using Metabond cement (Parkell, Inc).

In some optogenetic experiments, we targeted the dmPFC by implanting two 400 μm optic fibers 1.25 mm anterior from Bregma, 0.6 mm lateral (bilaterally). A small craniotomy (0.5-0.7 mm in diameter) was made and the tip of the fiber was placed on the brain surface. The fibers were positioned at a 5-8° angle from the sagittal plane (so that both fibers were in the coronal plane) to provide enough space for connecting two patch cables during optogenetic experiments.

In other optogenetic experiments, we targeted the anterior part of the dentate (lateral) nucleus (5.8 mm posterior, 2.2 mm lateral, 2.8 mm ventral relative to bregma). A small craniotomy (0.5-0.7 mm in diameter) was made 6.7 mm posterior, 2.2 mm lateral from bregma. Two 200 μm optic fibers were implanted bilaterally at 28° angle from the coronal plane (so each fiber was in a parasagittal plane), to avoid damage to the transverse sinus. An incision was made in dura and the fiber was inserted and lowered 1.8 mm into the brain (10 μm/s).

Black dental cement (Lang Dental, 1503BLK) was applied around the fiber to block laser light during the experiments, and dental cement (Bosworth Fastray Pink, Keystone Industries) was applied around it to reinforce the implant.

### Behavioral Training

During experiments, the headplate was attached to a pair of machined rods via 2–56 machine screws and the mouse was free to walk on top of the freely rotating foam cylinder (cylindrical treadmill). A high-speed (200 frames/s) monochrome camera (Allied Vision) and infrared illumination directed at the right side of the mouse’s face were mounted on a separate post via knuckle joints. Additional posts and clamps were used to hold a plastic tube for airpuff delivery, and an LED. The airpuff duration was 10-30 ms (20–30 PSI source pressure set in puffer). The stream of air was directed to the cornea via a 23 gauge blunt tip needle placed 3-5 mm from the mouse’s right eye. These pulse durations resulted in 6–8 PSI puff intensities measured at the end of the plastic tube leading to the needle. The pressure and duration of the periocular airpuff were set for each mouse to produce a full reflexive blink when delivered alone. All stimuli were controlled by an Arduino Due microcontroller.

Mice were habituated to head-restraint for two days prior to beginning the training sessions by placing them on top of the cylinder with their heads fixed for 30-60 minutes. No stimuli were delivered during this time. We then began training sessions in one of the following 3 behavioral tasks, typically a single session per day comprising approximately 100 trials (intertrial interval: 15-20 seconds).

#### I. The Omitted Oddball task

In each trial the mouse was presented with a train of flashes followed by an airpuff to the eye. Each flash was an LED (Luxeon Star) pulse of green light. The LED was placed above the animal’s head. The number of flashes in the train was randomly selected each trial from a uniform distribution, between 10 and 15 (including both values). The duration of each flash was 35 ms. The interval between flashes (between the beginning of a flash and the beginning of the next flash) was kept constant: 300 ms (for most experiments, including all optogenetic experiments), 150 ms or 450 ms. The interval between the offset of the last flash and the onset of the airpuff was always 200 ms longer than the interval between the flashes, i.e. the airpuff was always delivered 200 ms after the time of the “omitted” flash stimulus. Masking blue light was delivered using a laser and a patch cable of the same type as the one used for optogenetic inhibition (473 nm, Blue Sky Research), placed 1-2 cm beyond the mouse’s head. The masking light was kept flashing throughout the entire training session with pulses of random duration (20 to 200 ms) and random intervals between them (50 to 300 ms).

For training mice in the omitted oddball task with periodic stimuli of other sensory modalities, we used 35 ms bursts of white noise (90 dB) as a sound stimulus, or an electric stimulation of the mouse’s tail (0.3 mA, train duration 35 ms, duty cycle 4-12%, adjusted to minimize startle response).

#### II. Trace conditioning

In each trial the mouse was presented with a single green light flash (duration 35 ms) as the conditional stimulus, followed by an airpuff to the eye as the unconditional stimulus. The inter-stimulus interval between the offset of the flash and the onset of the airpuff was 200 ms.

#### III. Trace offset conditioning

In each trial the mouse was presented with a single green pulse of light of random duration (3-4.5 sec) as the conditional stimulus, followed by an airpuff to the eye as the unconditional stimulus. The inter-stimulus interval between the offset of the light pulse and the onset of the airpuff was 500 ms.

### Optogenetic inhibition

We used opto-blinks, brief laser pulses of 35 ms duration, to inhibit neural activity with high temporal precision at different moments during the training trials. The activity of dentate nucleus neurons was transiently suppressed by photo-activating the axon terminals of ChETA-expressing GABAergic Purkinje cells in L7-ChETA mice. We used blue (473 nm, Blue Sky Research) laser light delivered through optical fibers with 200 μm core diameter. Laser powers were in the range of 40 to 80 mW, measured at the tip of the patch cable. The activity of dmPFC neurons was transiently suppressed by photo-activating ChR2-expressing parvalbumin GABAergic interneurons in PV-IRES-Cre x Ai32 mice, using blue (473 nm) laser light delivered through two 400 μm optical fibers implanted bilaterally on the surface of the brain. Laser powers were in the range of 1.5-2 mW, measured at the tip of the patch cable. The patch cables and implants were wrapped around with black tape during the experiments.

### Histology

After mice with dentate nucleus implants completed the behavioral experiments, marking lesions were made via intense light stimulation through the optical fibers (Shanghai Laser & Optics Century Co., BL473T3–150FC, 60 mW for 40 s). Three days later, mice were euthanized and brains were extracted. Tissue was stored at 4 degrees C in 4% paraformaldehyde (Thermo Scientific Chemicals, AAJ19943K2) overnight before being transferred to a 30% sucrose solution in PBS for 3–5 days. Tissue was sectioned at 50 μm thickness on a microtome (Thermo Scientific, Microm HM 450) and stained with toluidine blue. Photomicrographs were captured using a Zeiss microscopy setup (Axio Imager microscope and ZEN software, blue edition).

### Data analysis

Data were analyzed in Matlab using custom written software. Eyelid movements were measured frame-by-frame from the high-speed videos by calculating the area of the eyelid visible within an elliptical region of interest. Raw pixel counts were normalized into units of fraction eyelid closure (FEC), which ranges from zero (fully open) to one (fully closed). Eyelid traces for each trial were extracted from the video frames.

Movement initiation latency was determined using an algorithm designed to identify sustained eyelid-closing movements while excluding small or transient fluctuations and preserving the earliest detectable phase of the movement. Eyelid movement traces were first converted to frame-by-frame velocity (change in FEC per 5-ms frame). For each frame, mean velocity was computed over a sliding 15-ms window (three frames), and frames where this mean exceeded 0.5 FEC/s were marked as above-threshold. Candidate movements were then identified as epochs in which at least 25 ms (five consecutive frames) remained continuously above threshold. Candidate movements were considered valid if the total change in FEC across the movement exceeded 0.1. For movements meeting this criterion, onset was defined as the first frame of the candidate movement.

All time scales in the paper take the moment of stimulus omission as zero, and movement initiation latencies are calculated relative to the moment of stimulus omission. The moment of stimulus omission equals the onset time of the last stimulus plus the stimulus period.

### Computational model of movement initiation

#### I. Movement initiation without optogenetic inhibition

To model the onset time for movement initiation in the omitted oddball task, we assumed that the latency of the ‘go’ signal in the dmPFC varies across trials as a normally distributed variable with mean ***m_g_*** and standard deviation ***s_g_***. Once the ‘go’ signal arrives in the cerebellum, the activity of the dentate nucleus starts to increase linearly from baseline ***B***, at a rate ***r***, until it reaches a threshold ***T***, when it triggers movement initiation. The rate ***r*** is a normally distributed variable with mean ***m_r_*** and standard deviation ***s_r_***. Ramp duration is (**T**-**B**)/**r**, and ***B*** and ***T*** are arbitrary values whose difference ***T***-***B*** is a scale factor that defines the units for ***m_r_*** and ***s_r_***. ***B*** was set to 10 a.u. (arbitrary units) and ***T*** was set to 20 a.u. Movement initiation latency is the sum of the ‘go’ signal latency and the ramp duration. The model is described by four parameters (***m_g_***, ***s_g_***, ***m_r_***, ***s_r_***).

The model assumes that the average time of the ‘go’ signal, in expert mice, is the moment of stimulus omission because this is the earliest point in time when the animal is able, in principle, to detect the omission and initiate the movement. Thus, our estimate of ***m_g_*** was 0 ms.

The probability of ‘go’ signal generation was calculated as a frequency of movements in control trials (without laser), which was 0.57 in the mice with dmPFC fiber implant, and 0.58 in the mice with the dentate fiber implant.

To estimate free parameters ***s_g_***, ***m_r_***, ***s_r_*** we used grid search with 3D parameter sweep (***s_g_*** and ***s_r_*** values were shared between all mice, ***m_r_*** values were mouse-specific). For each experimental session we calculated the histograms of observed and simulated latencies (histogram time bin 10 ms, 1000 simulated trials per session) and calculated the sum of squared deviations between observed and simulated movement initiation probabilities across all time bins. Simulated trials with negative values of ***r*** were excluded (∼0.5% of trials). Optimal parameter values identified using this approach were ***s_g_*** = 38.5 ms, ***m_r_*** = 0.090 a.u./ms average across mice (range 0.04-0.12 a.u./ms), ***s_r_*** = 0.035 a.u./ms.

#### II. Optogenetic inhibition of dmPFC

To model the effects of optogenetic inhibition of dmPFC we introduced one additional free parameter, ***u_PFC_***, which corresponds to the duration of dmPFC inhibition (may differ from the duration of laser pulse). If the ‘go’ signal is generated while the dmPFC is inhibited, then the signal will be eliminated and the ramp of activity in the cerebellar dentate nucleus will not start.

This parameter was optimized using 1D sweep, while other parameters (***s_g_***, ***m_r_***, ***s_r_***) were fixed at the optimal values. The cost function used for this optimization step was the sum of squared differences between observed and predicted probabilities of movement initiation in the dmPFC optogenetic experiments, averaged across mice and normalized by probabilities of movement initiation in trials without laser. The optimal value of ***u_PFC_*** was 93 ms.

#### III. Optogenetic inhibition of dentate

To model the effects of optogenetic inhibition of the cerebellar dentate we used two additional free parameters: ***u_LN_***, duration of dentate inhibition (may differ from the duration of laser pulse) and ***f_ramp_***, a multiplier that changes the ramping rate ***r*** as described below.

In the model, dentate inhibition affects movement initiation according to the following 3 rules:

1. The ramping cannot start during inhibition.
2. If the ‘go’ signal has arrived at the dentate during inhibition then the ramp starts from the baseline immediately after the inhibition ends. The slope of the ramp will be ***r_new_*** = ***f_ramp_*** \****r***.
3. If the inhibition starts during an ongoing ramp (before dentate activity reaches threshold ***T***) then the ramp stops. After the end of inhibition, the ramp resumes from the activity level where it was halted by the inhibition. The slope of the ramp will be ***r_new_*** = ***f_ramp_*** \****r***.

These two parameters (***u_LN_*** and ***f_ramp_***) were optimized using 2D sweep, while other parameters (***s_g_***, ***m_r_***, ***s_r_***) were fixed at the optimal values. The cost function used for this optimization step was the sum of squared differences between observed and predicted delays of movement initiation in the dentate optogenetic experiments, averaged across mice. The optimal value of ***u_LN_*** was 107 ms and the optimal value of ***f_ramp_*** was 2.54.

### Statistics and reproducibility

Tests were performed in MATLAB (MathWorks, version R2025b). No statistical methods were used to pre-determine sample sizes, but our sample sizes are similar to those reported in previous publications. All statistical tests were two-tailed. Comparisons between two paired groups (e.g., laser versus no-laser conditions within the same mice) were performed using paired t-tests. Comparisons between two independent groups were performed using two-sample t-tests. One-sample tests against a null value were performed using one-sample t-tests. Comparisons across three groups (e.g., movement latencies across the three stimulus period conditions) were performed using one-way ANOVA. When multiple time points were tested within the same dataset, family-wise alpha was maintained at 0.05 using the Bonferroni correction. Exact P values, test statistics, and degrees of freedom are reported in the figure legends. Error bars represent SEM unless otherwise noted. Eyelid trace averages in figures show 95% confidence intervals.

No data were excluded from the behavioral analyses. For the dentate nucleus optogenetic experiments, 2 mice were excluded from the dataset because of incorrect fiber targeting. Experiments were not independently replicated in separate cohorts and data collection and analysis were not performed blind to the conditions of the experiments. However, all key comparisons were made within animals (laser vs. no-laser trials interleaved within the same sessions), and results were consistent across mice within each group. Each mouse was tested in multiple sessions.

## ACKNOWLEDGMENTS

Supported by grants to JFM from the National Institutes of Health (R01MH093727; R01NS142192). Drawing of experimental setup in Figure 1A by A. Assie.

## AUTHOR CONTRIBUTIONS

M.M, G.W & J.M. designed the experiments. J.M. supervised the project. G.W. conducted the control experiments involving Trace Offset conditioning and collected preliminary data to help establish the Omitted Oddball task. M.M conducted all the remaining behavioral and optogenetics experiments. M.M analyzed and curated all original data collected for this publication. M.M made all figures. M.M & J.M. wrote the original draft of the paper.

---

**Extended Data Fig. 1.**
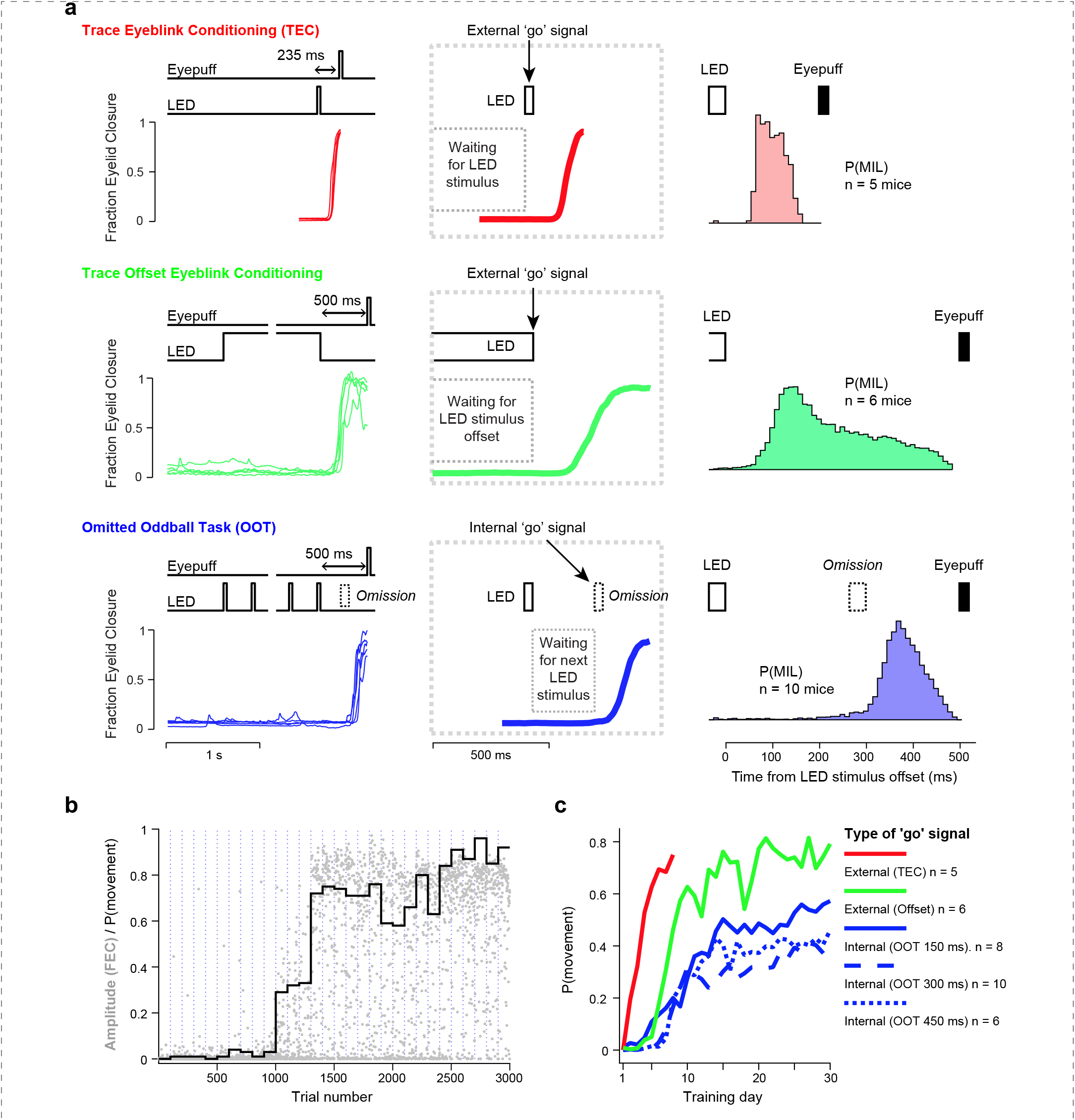
Comparison of performance in tasks with internal and external ‘go’ signals. In order to confirm that ‘go’ signal is generated internally in the Omitted Oddball Task (OOT), we compared distributions of movement initiation latencies in OOT and two tasks with external ‘go’ signals. In one task, a single LED flash served as ‘go’ signal (Trace Eyeblink Conditioning, TEC). In another task the offset of a long, variable duration, LED stimulus served as ‘go’ signal (Trace Offset Eyeblink Conditioning). To enable side-by-side comparison with OOT, the duration of this long stimulus was the same as duration of the train of flashes in OOT (3-4.5 s). Also, the interval between long stimulus offset and the eyepuff was the same as the interval between the last LED flash offset and the eyepuff in OOT (500 ms). **a,** Schematics of three tasks and characteristic movement latencies. Left: timing of stimuli and example eyelid movements Middle: relationship between stimuli and ‘go’ signals. Right: probability distributions of movement initiation latencies (MIL), relative to last LED stimulus offset. In TEC and Offset tasks the movements start soon after external ‘go’ signal, while in OOT movements start after LED stimulus Omission, when no external cues are present. **b,** Learning curve for one example mouse trained with OOT. Gray dots show movement amplitudes in each trial. Movement amplitude is the fraction eyelid closure (FEC) at the moment the eyepuff is delivered. Black staircase plot, probability of movement with amplitude above the threshold of 0.1 FEC, for each training session. Vertical dotted lines indicate start of each session. Several days of training were required before the mouse could start making correct movements. **c,** Averaged learning curves. Y-axis shows averaged (across mice) probabilities of movement with amplitude above the threshold of 0.1 FEC. The Omitted Oddball Task was more challenging for animals to learn, compared to tasks with external ‘go’ signals.

**Extended Data Fig. 2.**
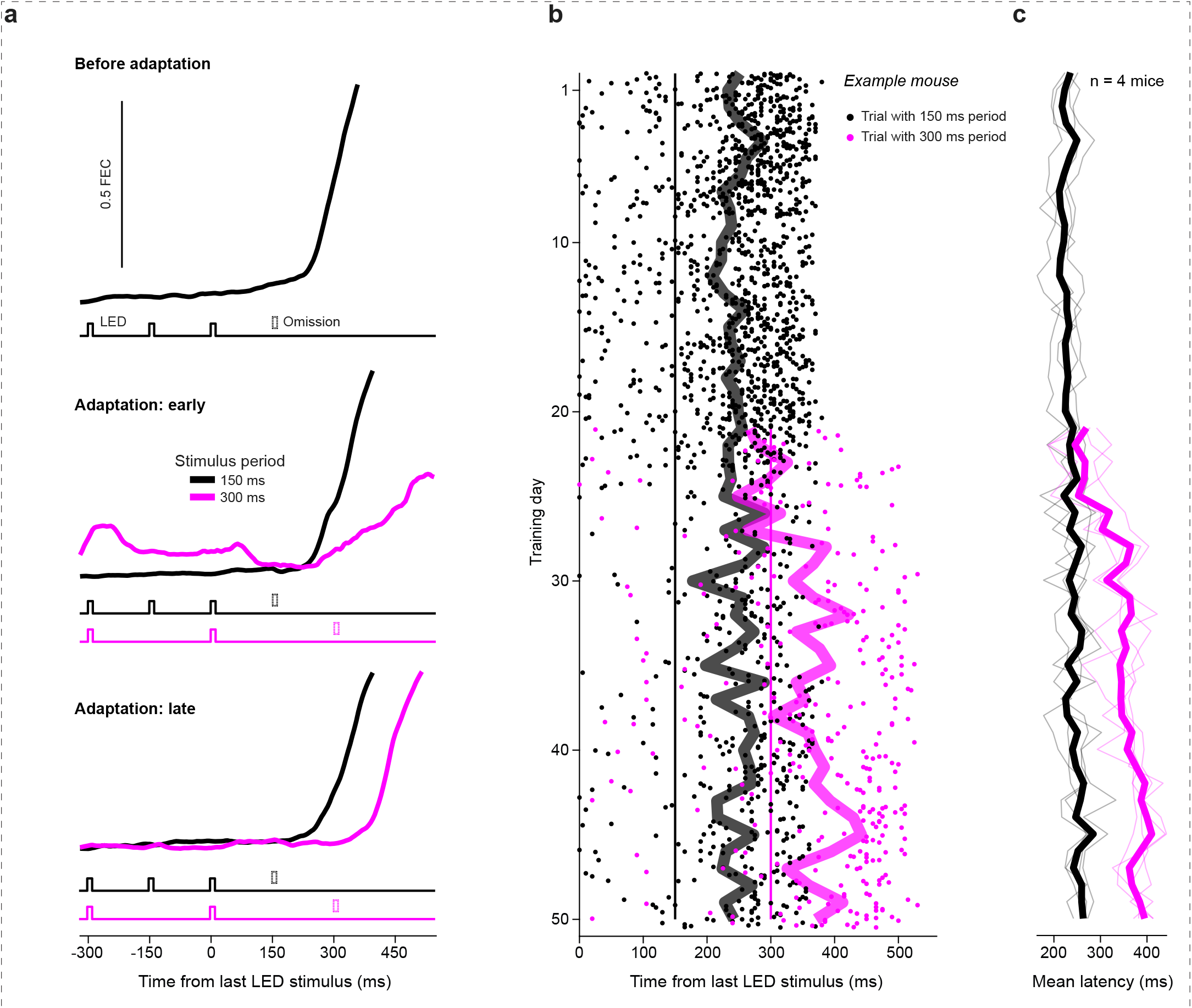
Adaptive timing of internally-driven movements. The Omitted Oddball Task requires animals to keep track of time passed after last LED stimulus and initiate the movement when the time exceeds LED stimulus period. We asked if mice were able to adjust movement initiation latency when the period changed. Mice were initially trained with 150 ms period. When performance stabilized, mice were trained with a mix of 150 ms period trials and 300 ms period trials (80% and 20% in first 3 days, 50% and 50% later on), **a,** Averaged eyelid movement traces aligned to last LED stimulus, in trials with 150 ms period (black) and 300 ms period (magenta), before and during the adaptation. An example mouse. **b,** Movement initiation latencies during initial training with 150 ms period and adaptation to new 300 ms period. Each marker shows one trial. Thick lines show averaged latencies across trials. Same example mouse as in **(a)**. After few days of adaptation the mouse was able to adjust movement initiation latency in each trial, depending on what the LED stimulus period was in this particular trial. Vertical lines indicate the moment of stimulus omission for each trial type. **c,** Averaged movement initiation latencies relative to last LED stimulus, during initial training and during the adaptation (n = 4 mice). Thin lines, averages across trials for each mouse. Thick lines, average across mice. All mice were able to adjust the movement initiation latency adaptively.

**Extended Data Fig. 3.**
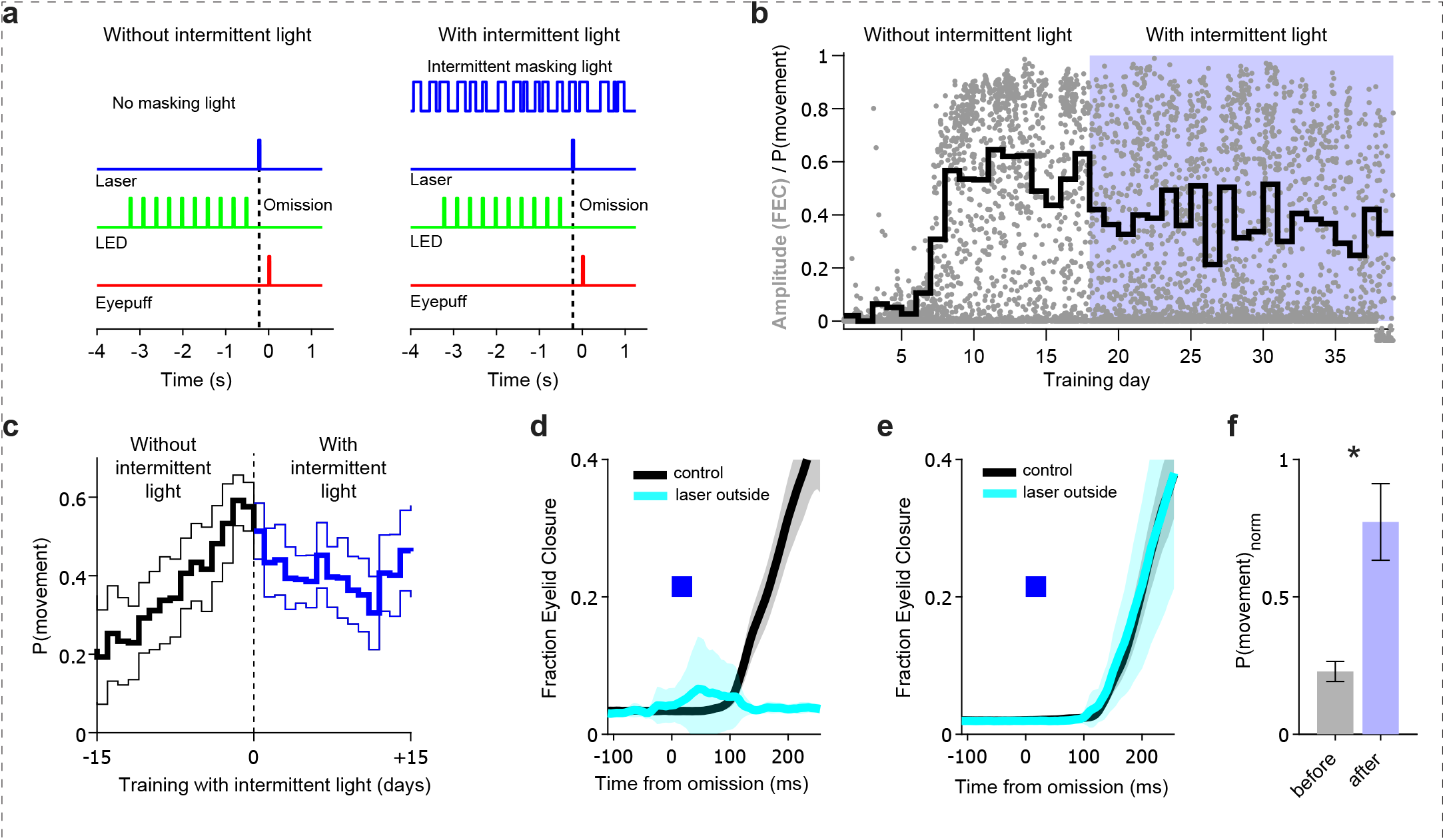
Using intermittent background light for masking. The laser pulse used for optogenetic inhibition can be seen by the mouse and needs to be masked. We used an intermittently flashing blue light for masking. The light was delivered by another laser (same wavelength and power), via a patch cable placed beyond the head of the animal, but not connected to the implant. It emitted a random sequence of flashes with variable duration. **a,** Schematic representation of stimuli delivery with and without intermittent blue light flashes in the background. **b,** Learning performance in the presence of intermittent light in one example mouse. The intermittent light was introduced on day 18. Gray dots show movement amplitudes in each trial. Movement amplitude is the fraction eyelid closure (FEC) at the moment the eyepuff is delivered. Black staircase plot, probability of movement with amplitude above the threshold of 0.1 FEC, for each training session. Although performance was slightly lower in the presence of intermittent light, the mouse was still able to initiate movements correctly in many trials. **c,** Averaged learning curves before (black) and after (blue) addition of intermittent light, n = 18 mice. Thin lines show 95% CI. The intermittent light decreased performance, but it remained sufficient to perform optogenetic experiments. **d,** An example session demonstrating necessity of masking. The laser pulse was delivered at the time of the LED stimulus omission, while the laser was not connected to the implant (’laser outside’). Shaded areas show 95%CI. The experiment was done before training the mouse with intermittent light. The laser pulse almost completely eliminated movements, even though no optogenetic inhibition could happen. Therefore, the laser pulse should be masked in order to probe the effects of optogenetic inhibition. **e,** same as in **(d)**, but after training the same mouse with intermittent light in the background. Now, the mouse was able to initiate the movements. **f,** Effects of laser pulse before and after training with intermittent light. The pulse was delivered from the laser not connected to the implant, at the time of the LED stimulus omission. Normalized P(movement) is a ratio of movement probabilities in trials with and without laser pulse. Error bars show SEM. n = 8 mice. Paired t-test, t(7) = −3.47, P = 0.010. After training mice with intermittent background light, the disruptive effect of external laser illumination was markedly reduced, allowing mice to reliably initiate movements despite delivery of the laser pulse at the time of the omitted stimulus.

**Extended Data Fig. 4.**
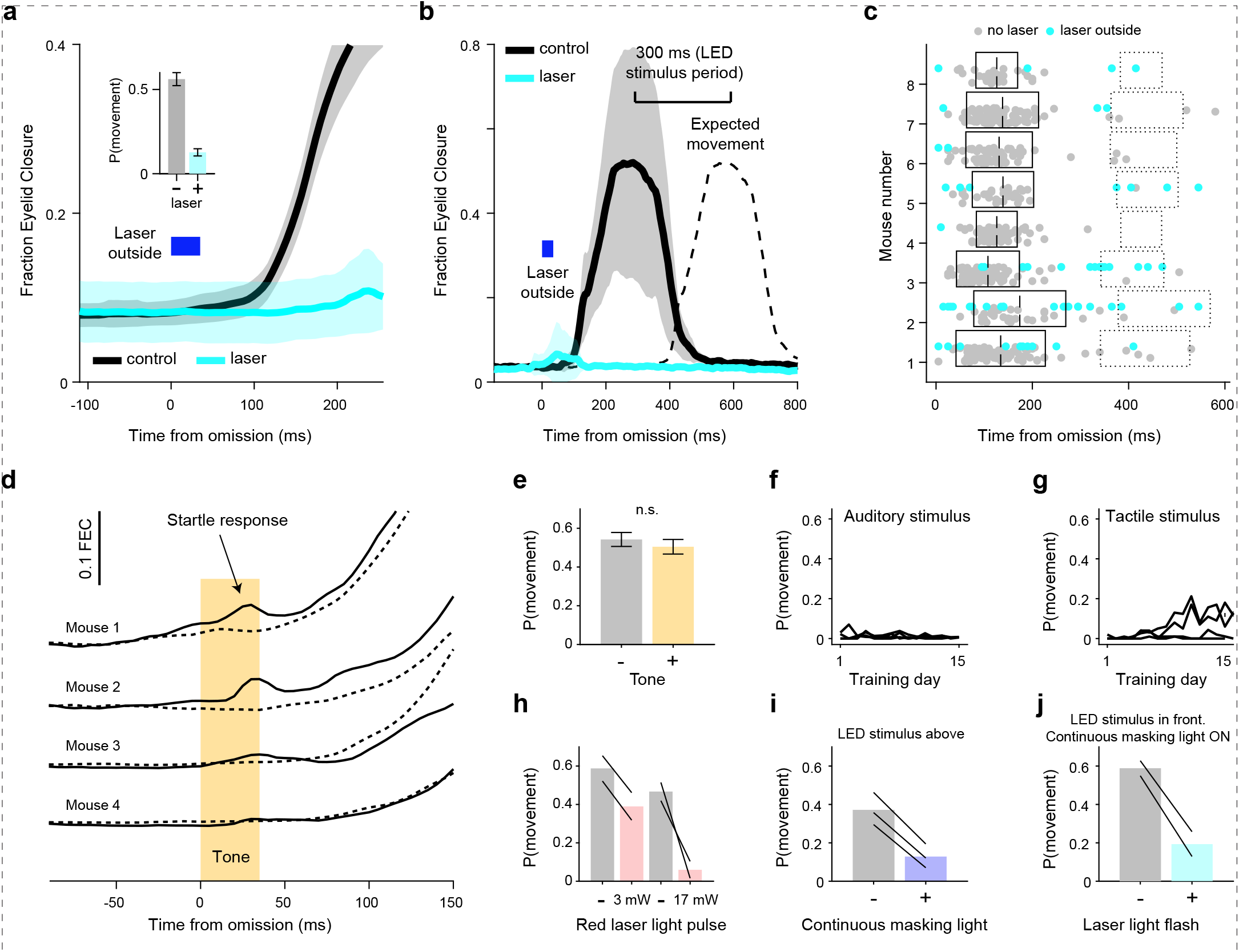
Confounding effects of laser light and assessment of alternative approaches to masking. Because the periodic stimulus used in the Omitted Oddball Task (OOT) was visual, any visible light associated with optogenetic perturbations could provide an additional sensory cue and confound the interpretation of behavioral effects. We therefore tested whether externally visible laser illumination, delivered without coupling the laser to the cranial implant, affected task performance, and evaluated alternative strategies for masking this visual artifact. **a,** An example session with laser pulse delivered at the time of the LED stimulus omission. The laser was not connected to the implant (’laser outside’). The experiment was done before training the mouse with intermittent light. Shaded areas show 95%CI. *Inset*: n = 8 mice. Paired t-test, t(7) = 11.15, P < 0.001. **b, c**, If visible laser light was perceived as an additional stimulus in the periodic LED train, mice would be expected to delay movement initiation by approximately one stimulus period. **b**, Example session showing movement initiation latencies on trials with and without laser illumination delivered outside the cranial implant. **c**, Movement initiation latencies across n = 8 mice tested before training with intermittent masking light. Dots show individual trials with laser illumination outside the implant (blue) and control trials without laser illumination (gray). Solid black rectangles indicate the mean ±SD latency for control trials. Dotted rectangles indicate the latency window expected if mice treated the visible laser pulse as an additional LED stimulus. Delayed movements in this window were rare, suggesting that visible laser illumination was not perceived as an extra stimulus in the LED sequence. **d,** The laser pulse could eliminate the movements if the animals were especially susceptible to interference from any unexpected sensory input during LED stimulus omission. But when we delivered an auditory tone (10 kHz, 35 ms) instead of laser pulse the performance was not impaired, even though the mice made small startle responses to the tone. Dashed traces show averaged movements in trials without tone. **e,** Movement initiation probability did not change when the tone was played at the time of the LED stimulus omission. Error bars show SEM. n = 4 mice. Paired t-test, t(3) = −1.67, P = 0.193. This observation suggests that we could reduce the impairment caused by the laser pulse if we were able to train the mice with a periodic stimulus of a different sensory modality (other than visual). **f,** However, training mice with bursts of white noise instead of LED flashes was unsuccessful (stimulus duration 35 ms, stimulus period 300 ms, n = 4 mice). **g,** Training with mild electric stimulation of the tail was not successful either (stimulus duration 35 ms, stimulus period 300 ms, n = 4 mice). Therefore, we continued using visual periodic stimulus (LED). **h–j,** Evaluation of alternative strategies to reduce the visual saliency of laser illumination. **h**, Replacing the blue laser (473 nm) with a red laser (635 nm) did not eliminate the behavioral effects of visible laser illumination, as movement initiation remained impaired during laser-outside trials (n = 2 mice). **i**, Continuous blue masking light was tested as an alternative to intermittent masking, but substantially impaired task performance (n = 3 mice). **j**, To improve performance under continuous masking conditions, the periodic LED stimulus was relocated from above the animal to a position directly in front of the mouse and its intensity was adjusted to increase its saliency relative to the background illumination. Although these modifications restored task performance (gray bar), laser-outside stimulation continued to disrupt movement initiation (cyan bar) despite the presence of continuous masking light of the same color (n = 2 mice). Based on these experiments, intermittent masking light was used in all subsequent studies.

**Extended Data Fig. 5.**
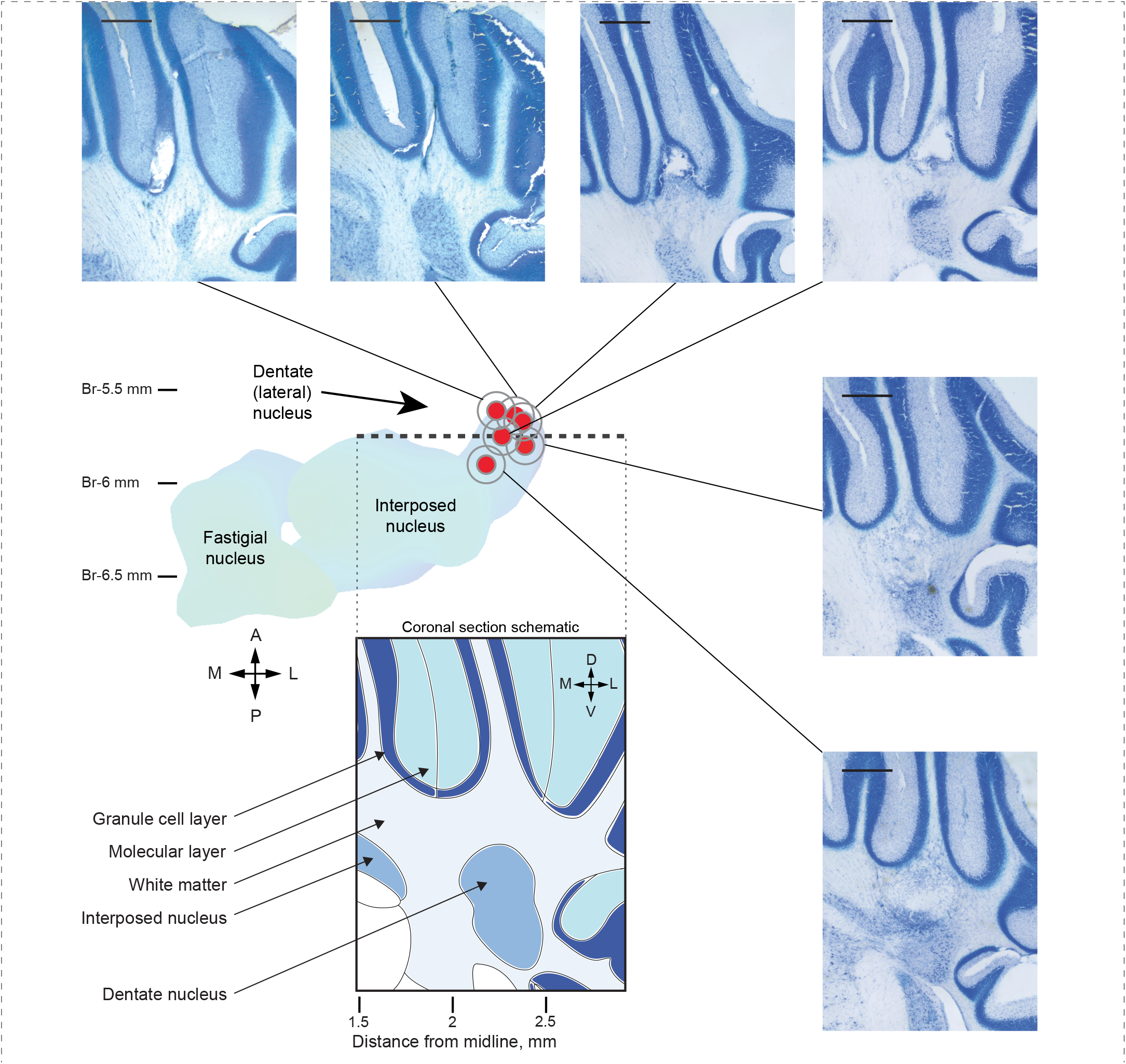
Locations of optic fibers in cerebellum. After finishing optogenetic experiments we performed histological examination to confirm the location of optic fibers above dentate nucleus. Thermal marking lesions were made using intense light stimulation through the optic fibers. Fiber locations were estimated by visual inspection of Nissl-stained coronal brain sections (50 μm thick) The schematic at the center shows top view of deep cerebellar nuclei. The gray outline around each circle shows optic fiber diameter (200 μm). Red circle, fiber center. Scale bars on coronal sections are 500 μm. FN, fastigial nucleus. IP, interposed nucleus. In all mice (n = 6) the fiber tips were in close vicinity of Dentate nucleus.

**Extended Data Fig. 6.**
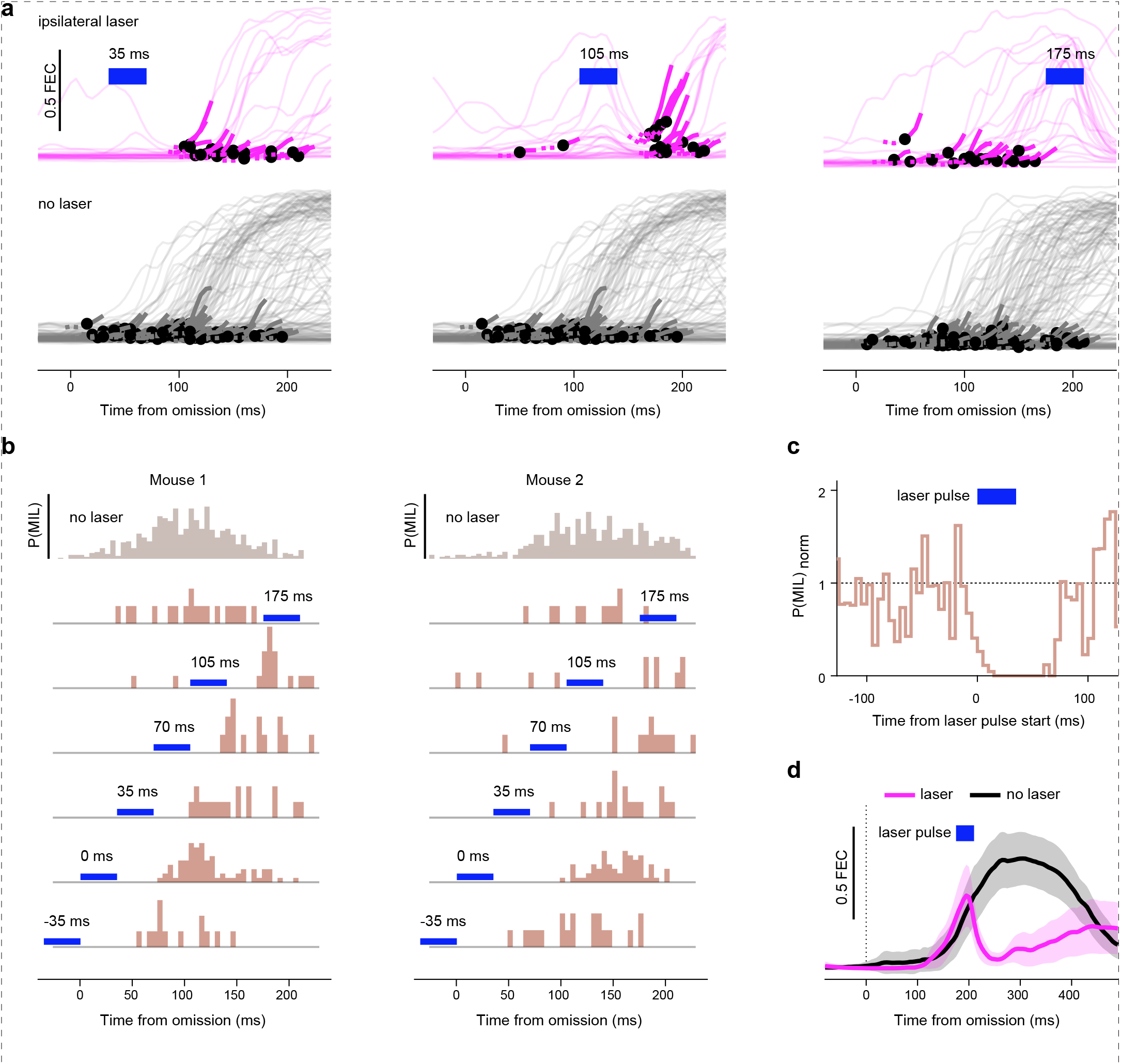
Dentate activity is necessary for movement initiation and execution. To investigate the reasons for movement initiation delay after dentate inhibition we inspected distributions of movement initiation latencies in trials with laser pulses. **a,** Example sessions with inhibition of dentate (same mouse, different laser pulse timing). Blue rectangles are laser pulses, text labels above them show pulse start time relative to LED stimulus omission. Black dots are movement onsets. FEC, fraction of eyelid closure. Movements did not start during laser pulse or shortly after it. **b,** Histograms of movement initiation latencies in control trials (top) or in trials with laser delivered to dentate at various time points (bottom), in two example mice. Blue rectangles are laser pulses. P(MIL), probability of movement initiation latency. The probability of movement initiation was nearly zero in a 60-70 ms window after laser onset. **c,** Histogram of movement initiation latencies aligned to laser pulse. The histogram bin heights (probability values) were averaged across multiple sessions where laser pulse was delivered at various time points relative to LED stimulus omission. In each session the bin heights from trials with laser were divided by corresponding bin heights in control trials without laser. n = 6 mice. This alignment and normalization of P(MIL) distributions allows us to pool data from multiple sessions and visualize the effect of laser pulse regardless of when it was delivered relative to LED stimulus omission. Movements were almost never initiated during the laser pulse or in the 30-40 ms window immediately after it. **d,** The effect of laser pulse, delivered to dentate, on the ongoing movements (example session). Traces show averaged eyelid movements across trials 95% CI (shaded area). Blue rectangle is the laser pulse. The inhibition stopped execution of ongoing movements.

**Extended Data Fig. 7.**
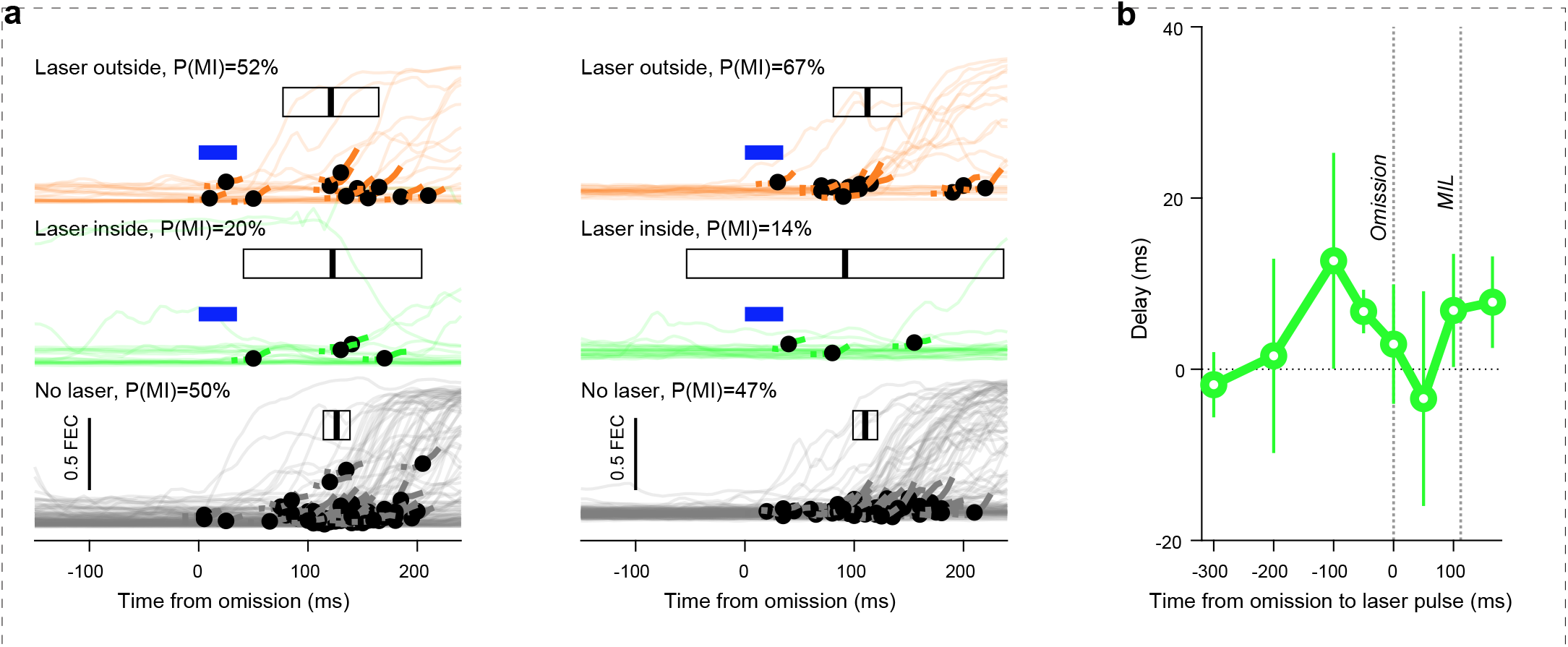
Inhibition of dmPFC does not change movement initiation latency. Since inhibition of Dentate delayed movements, we asked if inhibition of dmPFC has similar effect. **a,** Example sessions with dmPFC inhibition at the time of the LED stimulus omission in two different mice. Black dots show movement onsets in individual trials. Black rectangles show 95%CI of movement initiation latency, vertical bar inside the rectangle shows mean latency. P(MI), probability of movement initiation. Latencies did not change significantly, but the probability of movement initiation decreased. **b,** Averaged delays of movement initiation (differences between mean latencies in ‘laser inside’ and ‘laser outside’ conditions) in experiments with various laser timings. MIL, average movement initiation latency. Error bars show SEM. One-sample t-tests with Bonferroni correction (8 comparisons), n = 3 to 7 mice at each time point. No time points reached significance (all P > 0.05). Since the number of movements is strongly reduced after inhibition the test may lack statistical power to detect any changes of latency (in some sessions the movements were initiated only in a few trials). Nevertheless, the results suggest that dmPFC inhibition affects only movement probability but not its latency.

**Extended Data Fig. 8.**
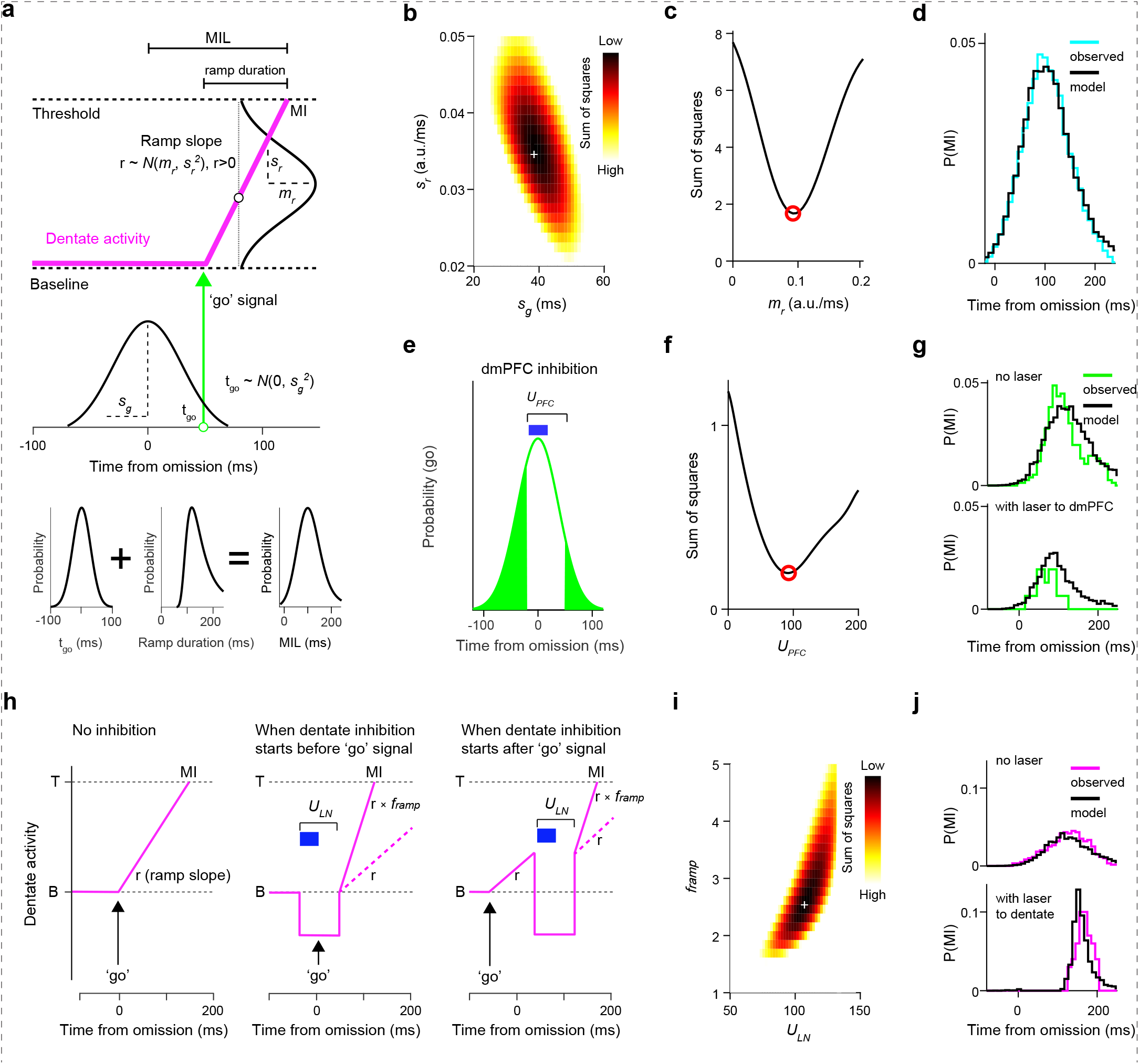
Computational modeling of movement initiation. We asked if the observed effects of optogenetic inhibition can be accounted for by a model based on hypothesized interactions between dmPFC and dentate (see Methods for details). **a,** Model schematic. The ‘go’ signal is generated in dmPFC and transmitted to Dentate, where it causes Dentate activity to ramp up. Once the activity reaches a threshold the movement is initiated. Both ‘go’ signal time and ramp slope are normally distributed variables. MIL, movement initiation latency, measured relative to the moment of LED stimulus omission. MIL is the sum of ‘go’ signal time and the duration of the ramp. *s_g_*, standard deviation of ‘go’ signal time. *m_r_*, mean slope of the ramp. *s_r_*, standard deviation of the slope. **b,** Estimation of parameters *s_r_* and *s_g_*, shared between all mice, used 2D parameter sweep. White cross symbol indicates the optimal pair of values. a.u., arbitrary units. **c,** Estimation of parameter *m_r_* in one example mouse. Red circle shows optimal value. a.u., arbitrary units. **d,** Observed and predicted movement initiation latencies in one example mouse (without optoge-netic inhibition, same mouse as in **(c)**). P(MI), probability of movement initiation. Time bin is 10 ms. **e,** A schematic of ‘go’ signal elimination by dmPFC inhibition. Parameter *U_PFC_* determines the duration of inhibition that starts with laser pulse (blue rectangle). White area under the green curves shows the part of ‘go’ signal probability distribution ‘carved out’ by the inhibition. **f,** Estimation of parameter *U_PFC_*. Red circle shows the optimal value. **g,** Observed and predicted movement initiation latencies after dmPFC inhibition at the moment of stimulus omission (example session). P(MI), probability of movement initiation. Time bin is 10 ms. **h,** A schematic showing effects of dentate inhibition on the ramping activity (see Methods for details). MI, movement initiation. T, threshold for movement initiation. B, baseline. r, ramp slope. *U_LN_*, the parameter that determines duration of inhibition. Blue rectangle, laser pulse. *f_ramp_*, the parameter that changes ramp slope after inhibition offset. Dashed magenta lines indicate how the ramp would continue if *f_ramp_* = 1. **i**, Estimation of parameters *U_LN_* and *f_ramp_* using 2D sweep. White cross symbol indicates optimal pair of parameters. **j,** Observed and predicted movement initiation latencies after dentate inhibition at the moment of stimulus omission (example session). P(MI), probability of movement initiation. Time bin is 10 ms.

